# Accurate protein structure determination from cryo-EM maps using deep learning and structure prediction

**DOI:** 10.1101/2025.10.23.684082

**Authors:** Tao Li, Ji Chen, Hao Li, Hong Cao, Sheng-You Huang

## Abstract

Cryo-electron microscopy (cryo-EM) has become the main-stream technique for macromolecular structure determination. However, due to intrinsic resolution heterogeneity, accurate modeling of all-atom structure from cryo-EM maps remains challenging even for maps at near-atomic resolutions. Addressing the challenge, we present EMProt, a fully automated method for accurate protein structure determination from cryo-EM maps by efficiently integrating map information and structure prediction with a three-track attention network. EMProt is extensively evaluated on a diverse test set of 177 experimental cryo-EM maps with up to 54 chains in a case at <4.0 Å resolutions, and compared with state-of-the-art methods including DeepMainmast, ModelAngelo, phenix.dock_and_rebuild, and AlphaFold3. It is shown that EMProt significantly outperforms the existing methods in recovering the protein structure and building the complete structure. In addition, the built models by EMProt also exhibit a high accuracy in model-to-map fit and structure validations. EMProt is freely available at http://huanglab.phys.hust.edu.cn/EMProt/.

## 1 Introduction

Proteins play important roles in many biological processes by themselves or interacting with other molecules. Therefore, determining the atomic structures of proteins in three dimensions is crucial for understanding molecular life processes. In the past decade, cryo-EM has significantly advanced the process to determine the structures of biological macromolecules, achieving the resolvability of individual atoms^1–3^. As a result, the Electron Microscopy Data Bank (EMDB) has seen an exponential increase in the number of deposited cryo-EM maps. Various methods have been developed for automated model building of cryo-EM maps^4–8^. Many macromolecular complex structures have been resolved and deposited in the Protein Data Bank (PDB)^9^. Nevertheless, model building of cryo-EM maps remains a labor-intensive and challenging work, and requires significant human expertise.

For years, a number of methods have been proposed to reduce the obstacles for accurately modeling structures from cryo-EM maps. Current cryo-EM model building methods can be broadly categorized into two types: *de novo* based modeling and template-based fitting. The *de novo* modeling methods, such as phenix.map to model^10^, MAINMAST^11^, DeepMM^12^, C-CNN^13^, A2-Net^14^, DeepTracer^15^, and Cryo2Struct^16^, often starts by detecting critical atom positions from map and threading them to chains. ModelAngelo^17^, as the state-of-the-art *de novo* modeling tool, has gained significant attentions. While these methods demonstrate the feasibility of modeling a structure directly from raw density maps without additional knowledge, they often fail in density regions with poor local qualities even for maps at near-atomic resolutions.

For template-based fitting approaches, researchers often fit a template structure into the density map as an initial placement and then refine the structure manually for multiple rounds. With the advancement of structure prediction algorithms like AlphaFold2 (AF2), fitting highly accurate predicted structures into cryo-EM maps is frequently adopted in the field. Many automated tools like gmfit^18^, Situs^19, 20^, EMBuild^21^, DEMO-EM^22, 23^, and phenix.dock and rebuild^24^ have been developed to minimize the need for manual intervention. These automated tools are capable of modeling a reliable structure if the predicted structure is sufficiently accurate to be fitted into the map. However, they often fail if the predicted global fold deviates significantly from the actual ground truth structure.

The difference between two modeling categories is sometimes blurred as several methods attempt to integrate the information from both the map and templates. CR-I-TASSER^25^ uses the C*α* atom position predicted from density map to re-select a suitable template, though it is primarily designed for segmented single-chain maps, and does not utilize structure predictions by AlphaFold^26, 27^. DeepMainmast^28^, an enhanced version of MAINMAST, shows the potential of automated AF2guided *de novo* modeling. However, the structure-map fitting in DeepMainmast is not exhaustive, and the integration of structure and map is also kind of empirical. As such, its practical applicability may be limited by time cost and modeling accuracy.

With the exponential increase of released cryo-EM maps at near-atomic resolutions, there is a pressing need for automated and accurate modeling for the complete structure from a map with weak local densities. Meeting the need, we introduce EMProt, an automated modeling approach for accurate protein structure determination from cryo-EM maps by effectively integrating *de novo* modeling and structure prediction using a three-track attention network. EMProt first utilizes deep leaning to predict the amino acid types and the positions of C*α* atoms from raw density maps. Then, a three-track attention network is used to build the all-atom structure by integrating the 1D and 2D density features of predicted C*α* positions and 3D backbone frame into reasoning of 3D atomic coordinates. Moreover, EMProt is also able to integrate predicted structures through a combination of structure alignment to *de novo*-built model and FFT-based structure-to-map fitting. EMProt is evaluated on a benchmark set of 177 experimental maps with resolutions better than 4 Å, and compared with state-of-the-art methods, including DeepMainmast, ModelAngelo, phenix.dock_and_rebuild (Phenix), and AlphaFold3 (AF3). Overall, EMProt achieves the best performance among the five approaches in terms of both recovering the protein structure and building the complete model.

## 2 Results

### 2.1 The workflow of EMProt

The overall workflow of EMProt is illustrated in Fig. 1a. EMProt first threads an initial protein model using only information from the density map. To achieve this, two separately trained Swin-Conv-UNet networks are employed simultaneously: one predicts the voxel-wise probabilities of N, C*α*, and C atoms, while the other predicts the amino acid type probability. Afterwards, the C*α* positions are derived from the probability map using a mean-shift algorithm. At near-atomic resolution, density maps often lack the precision needed to accurately detect all backbone atoms, let alone side-chain atoms. In addition, the predicted C*α* positions often deviate from their true locations. Therefore, to accurately build the all-atom structure, we design a three-track attention network ^26, 29^ to refine C*α* positions, and predict backbone frames and torsion angles from raw C*α* atoms and density map (Fig. 1c-d). The three-track attention network is trained by noising the PDB atoms and having the network to recover the true positions. As for feature generation, with initial C*α* positions, we extract the densities around each atom as 1D representation and the densities between pairwise atoms as 2D representation. The C*α* positions and randomly initialized rotation matrices are used as 3D backbone frames. We use self-attention and invariant point attention to update 1D representation with attention biases from the 2D representation and 3D backbone frames. We use axial-attention to update the 2D representation with attention biases from the 3D backbone frames. The trained network is able to model all-atom positions given only C*α* atoms by predicting backbone frames and torsion angles.

**Figure 1:**
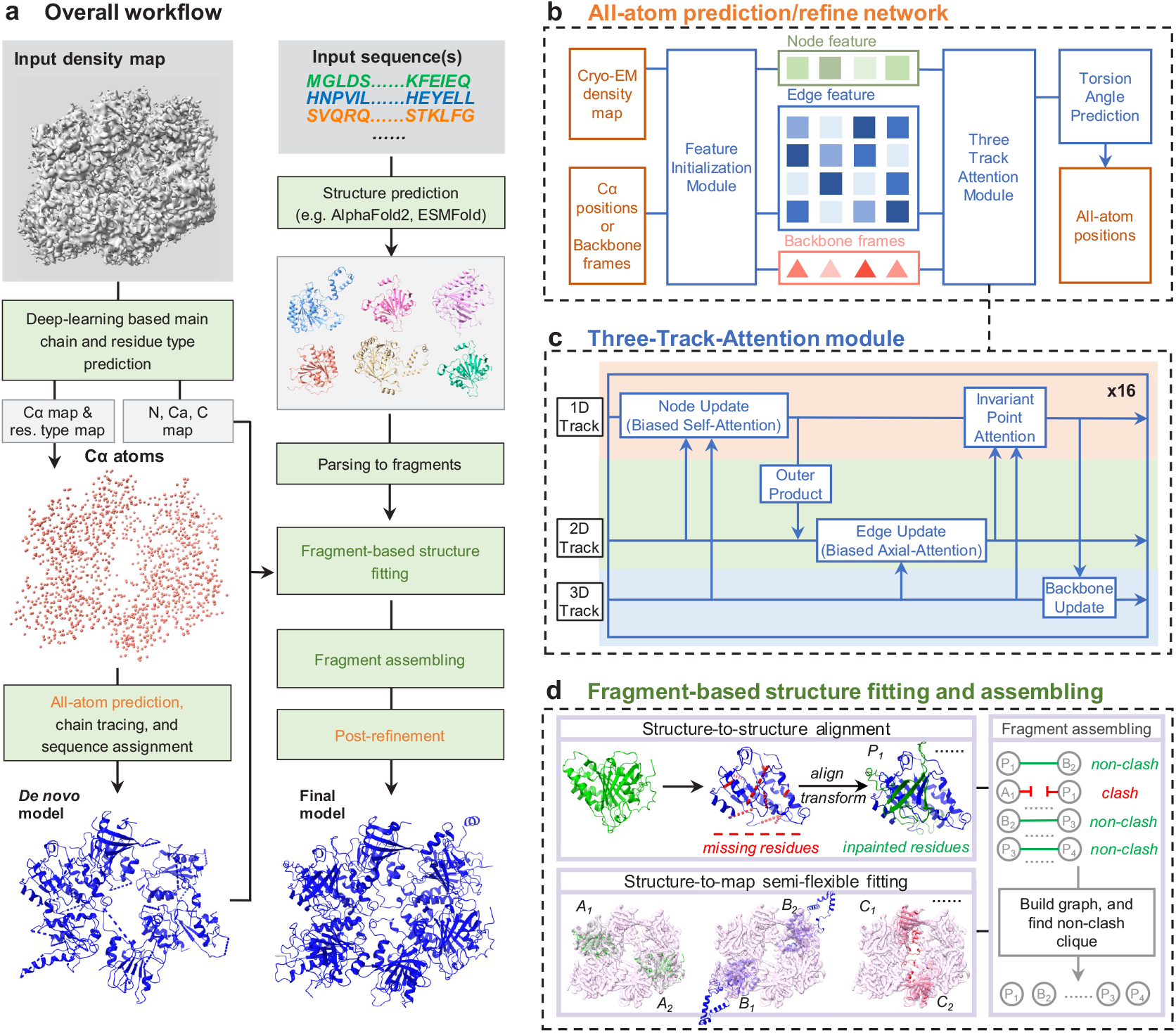
Overview of the EMProt framework. **a**, The workflow of EMProt. The input is the density map and sequence(s). In the *de novo* pipeline (left side), EMProt goes through C*α* prediction, backbone frame prediction, chain tracing and sequence assignment. In the integrative pipeline (right side), structure are predicted using input sequences and are fitting to the map in fragment level and output structure is assembled based on all fragments. Finally, a post refinement can be performed to further improve the model-to-map fit and structure quality of the built model. **b**, The all-atom prediction and refinement module. Node and edge features are initialized using C*α* atoms and density map. After the node, edge and backbone frames communicate through the Three-Track-Attention network, a final layer is used to predict torsion angles and all-atom positions are built accordingly. **c**, The Three-Track-Attention module. The biased self-attention is used to update node features, and the biased axial-attention is used to update edge features. The invariant point attention is used to fuse node, edge features and backbone frame information. **d**, The structure fitting and assembling module. The AF2 models are integrated by structure alignment and semi-flexible fitting. Finally, we select non-clash fragments on a graph built on all fragments.

After the backbone frames and torsion angles are predicted, we thread individual residues into continuous chains under constraints from peptide bond length. With the threaded chains, we assign the amino acid type probability for each path, and then perform an HMM search against the provided sequences^30, 31^. To achieve this, the initial amino acid probability for each residue is determined by finding the nearest voxel in the predicted amino acid type map. These probabilities are converted into the HMM profiles as position-specific scores using HMMER^17^, which are then used to search the input sequences and identify the matching regions, yielding the final residue sequence. With the amino acid identities assigned for each path, the side-chain atoms are constructed according to the predicted torsion angles.

After *de novo* modeling, EMProt may further integrate predicted structures by matching them with the density map (Fig. 1b). We use AF2 predicted structures in this study. Since the predicted model may have deviations from the true structure, EMProt will first split predicted chain models into fragments based on structural contacts. Then, EMProt aligns the AF2 fragments with the *de novo*-built fragments using TM-align^32^. The alignment establishes the residue correspondences between *de novo*-built fragments and AF2 structure fragments, enabling EMProt to: (i) Identify the regions in *de novo* fragments with incorrect amino acid identities, and (ii) Detect missing regions in the *de novo* model. If an AF2 fragment aligns well with a *de novo* fragment (e.g., with an RMSD *<* 1.5 Å), EMProt will refine the amino acid identities in the *de novo* model and fill in missing residues using the AF2-derived structure. Next, EMProt performs model-to-map fitting, enabling local sub-structural movements to improve the fit into the main-chain probability map through a semi-flexible refinement process. The corrected *de novo* fragments and the fitted AF2 structures are combined into a fragment library.

As multiple fragments may occupy adjacent positions, structural clashes can occur. To resolve this, EMProt applies a fragment-assembling algorithm that selects well-matched, non-clashing fragments to build the final model. A graph-based optimization algorithm is used, where each fragment is treated as a node and edges are added between non-clashing fragments. The algorithm solves a constraint optimization problem on this graph, retaining only the fragments that fit the density map while avoiding clashes. Finally, the assembled complex structure can be refined by a post-refinement program like phenix.real space refine^33^, which is recommended for improving the local geometry and model-to-map fit. In this study, all the models are refined by phenix.real space refine^33^ during the evaluations, unless otherwise specified.

### 2.2 Performance in recovering the protein structure

EMProt is first validated on a test set of 177 non-redundant experimental maps, and compared with state-of-the-art methods that employ different modeling strategies, including ModelAngelo (*de novo* modeling), Phenix (phenix.dock_and_rebuild, template-based modeling), and DeepMainmast (a hybrid of *de novo* and template-based modeling). Following previous studies^10, 15, 16^, three metrics, residue coverage, sequence match, and sequence recall, are used to evaluate the performance of different methods in recovering the protein structure. The corresponding values are calculated using phenix.chain comparison with a distance cutoff of 3.0 Å and C*α* atoms as the representatives^10^.

Figure 2 shows the bar-plots and head-to-head comparisons of the corresponding results by EMProt and the other three methods. As shown in Fig. 2a, d, EMProt outperforms the other methods in all metrics. Specifically, for residue coverage, EMProt achieves the highest average value of 91.20%, compared with 75.13% for DeepMainmast, 77.81% for ModelAngelo, and 79.25% for Phenix. Among the 177 maps, EMProt obtains a higher coverage than DeepMainmast for 130 cases, ModelAngelo for 163 cases, and Phenix for 131 cases. In addition, EMProt also performs better than the other methods in both sequence match and sequence recall, achieving the high values of 95.87% and 87.79%, respectively, compared with 83.80% and 68.72% for DeepMainmast, 94.98% and 75.08% for ModelAngelo, and 92.39% and 74.85% for Phenix (Fig. 2b-c, e-f). In details, EMProt has a better sequence match for 123, 111, and 83 cases and a better sequence recall for 136, 159, and 127 cases, compared with DeepMainmast, ModelAngelo, and Phenix, respectively. The sequence match metric only considers the built parts, whereas the sequence recall accounts for all residues in the PDB structure. In other words, both EMProt and ModelAngelo can correctly assign nearly all amino acid types in the built model, but EMProt correctly builds much more residues than ModelAngelo.

**Figure 2:**
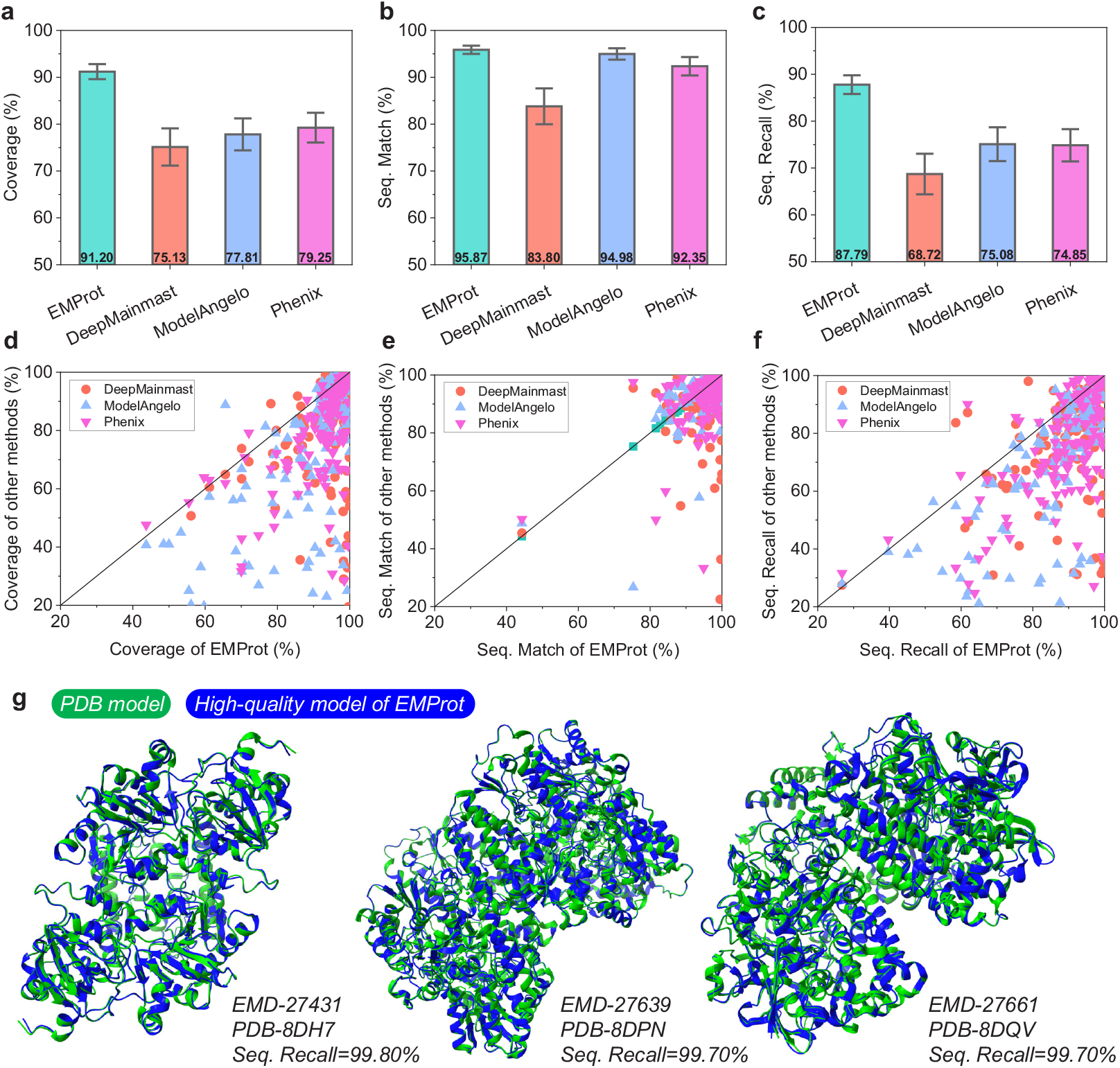
Comparison between EMProt, DeepMainmast, ModelAngelo, and Phenix in recovering the PDB structures on the test set of *n* = 177 maps. **a-c**, Bar plots of residue coverage, sequence match, and sequence recall. The bars indicate the means, and the error bars indicate 95% confidence interval. **d-f**, Head-to-head comparison of residue coverage, sequence match and sequence recall. **g**, Three examples of the high-quality models built by EMProt for three targets.

Figure 2g lists three EMProt-built models with *>*99% sequence recall. The targets are EMD-27431 (PDB: 8DH7), EMD-27639 (PDB: 8DPN) and EMD-27661 (PDB: 8DQV) from left to right, with resolutions of 2.99 Å, 2.49 Å, and 1.52 Å, respectively. It can be seen from the figure that the built models for three targets show a perfect overlap with the PDB structures even in loop regions. It is also noted that besides the showed three examples here, there are a total of 25 built models with a sequence recall of *>*99% and 100 models with sequence recall of *>*90%. Most of the cases have a resolution better than 3.0 Å, a relatively high resolution for EMProt to recover most of the residues from the map.

### 2.3 Examples in recovering the protein structure

Several examples are given in Fig. 3 to show how much the models built by EMProt, DeepMainmast, ModelAngelo, and Phenix recover a protein structure. Figure 3a shows the built models for target EMD-15220 (PDB: 8A7D) at 3.1 Å resolution. The built models are colored in blue, and the PDB structures are colored in green. For this target, EMProt achieves a sequence recall of 91.06%, which is significantly higher than 75.85% for DeepMainmast, 73.04% for ModelAngelo, and 76.68% for Phenix. When compared with the PDB structure, the EMProt model shows a good consistency with the reference PDB structure, while DeepMainmast and ModelAngelo fail to recover the residues in two regions, as indicated in the dashed rectangles, and Phenix fails in one of these regions.

**Figure 3:**
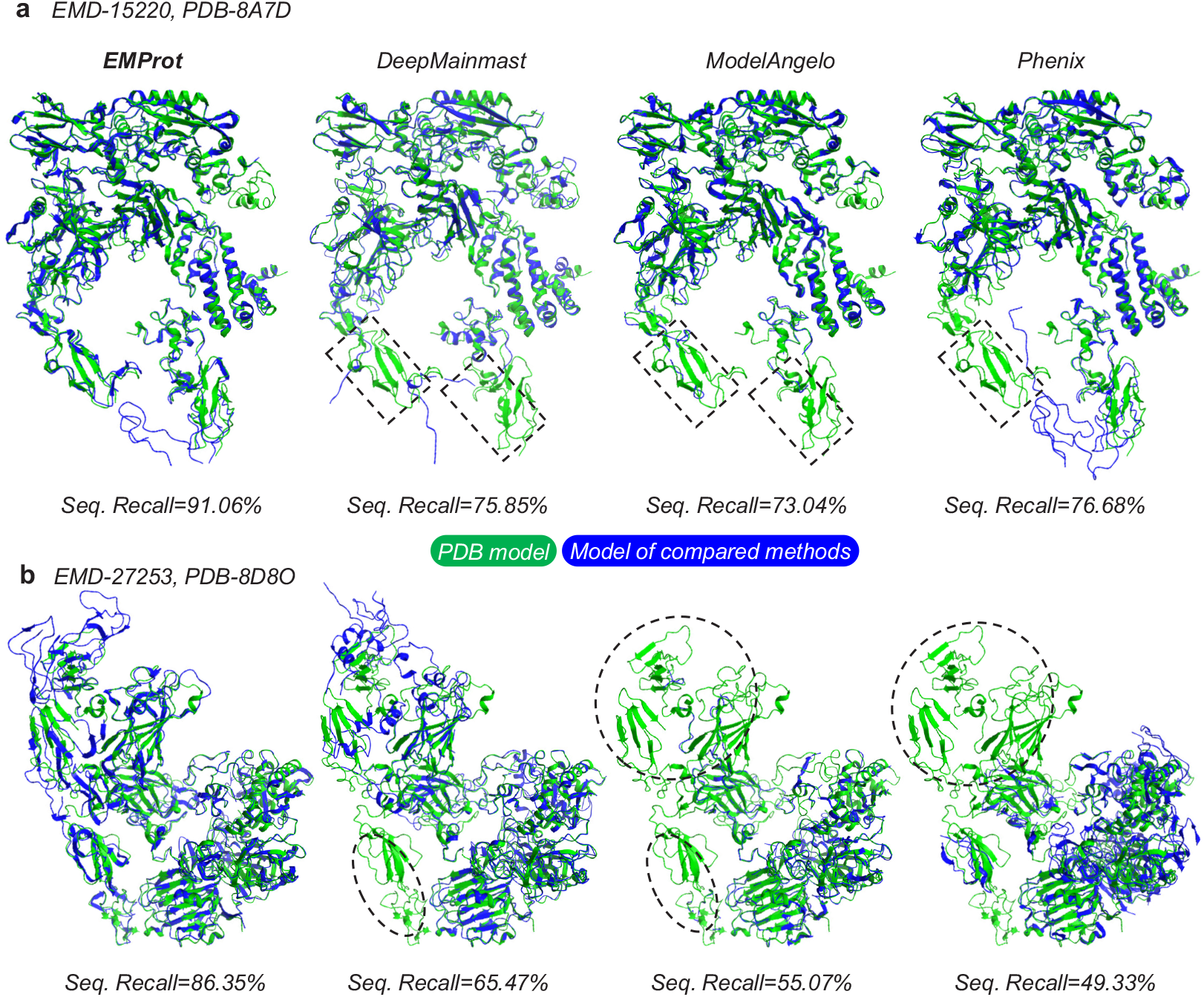
Comparison of the built models by EMProt, DeepMainmast, ModelAngelo, and Phenix in terms of recovering the PDB structures. **a**, Target EMD-15220 (PDB: 8A7D) at 3.1 Å resolution. **b**, Target EMD-27253 (PDB: 8D8O) at 3.4 Å resolution.

Figure 3b shows another comparison of the built models by four methods on target EMD-27253 (PDB: 8D8O) at 3.4 Å resolution. The sequence recall values are 86.35% for EMProt, 65.47% for DeepMainmast, 55.07% for ModelAngelo, and 49.33% for Phenix. Once again, EMProt demonstrates a superior performance by achieving the highest sequence recall, while the other three methods exhibit different degrees of structural deficiency and fail to build the structure in the regions, as indicated by dashed ellipses. Specifically, for ModelAngelo and Phenix, a very large area of the structure is missing, resulting a low sequence match of *<* 60%.

### 2.4 Performance in building the complete structure

We further evaluate the ability of EMProt and the other methods in building the complete structure using two global metrics: TM-score and RMSD. The metrics are calculated using USalign^34^ in multimer mode. Here, we intentionally exclude reporting the two metrics of ModelAngelo, as they are not suitable for the non-ordered structural fragments built by ModelAngelo. In addition, since protein multimer structure prediction is greatly improved in AF3, we also list its performance for comparison on this test set. The AF3 structures are predicted by the AF3 server and the one with the highest pTM score is selected out of five predictions.

Figure 4 shows the bar-plots and head-to-head comparisons of the corresponding results by EMProt and the other three methods (DeepMainmast, Phenix, and AF3). It can be seen from the figure that EMProt outperforms the other three methods in both TM-score and RMSD. For TM-score (Fig. 4a, c), EMProt achieves the highest average value of 0.90, compared with 0.69 for DeepMainmast, 0.82 for Phenix, and 0.84 for AF3. Among the 177 maps, EMProt obtains a higher TM-score than DeepMainmast for 132 cases, Phenix for 117 cases, and AF3 for 102 cases. Interestingly, AF3 performs well on this test set and even better than Phenix. Such a phenomenon can be understood because this test set is built before AF3 was released and some items are included in the training set of AF3.

**Figure 4:**
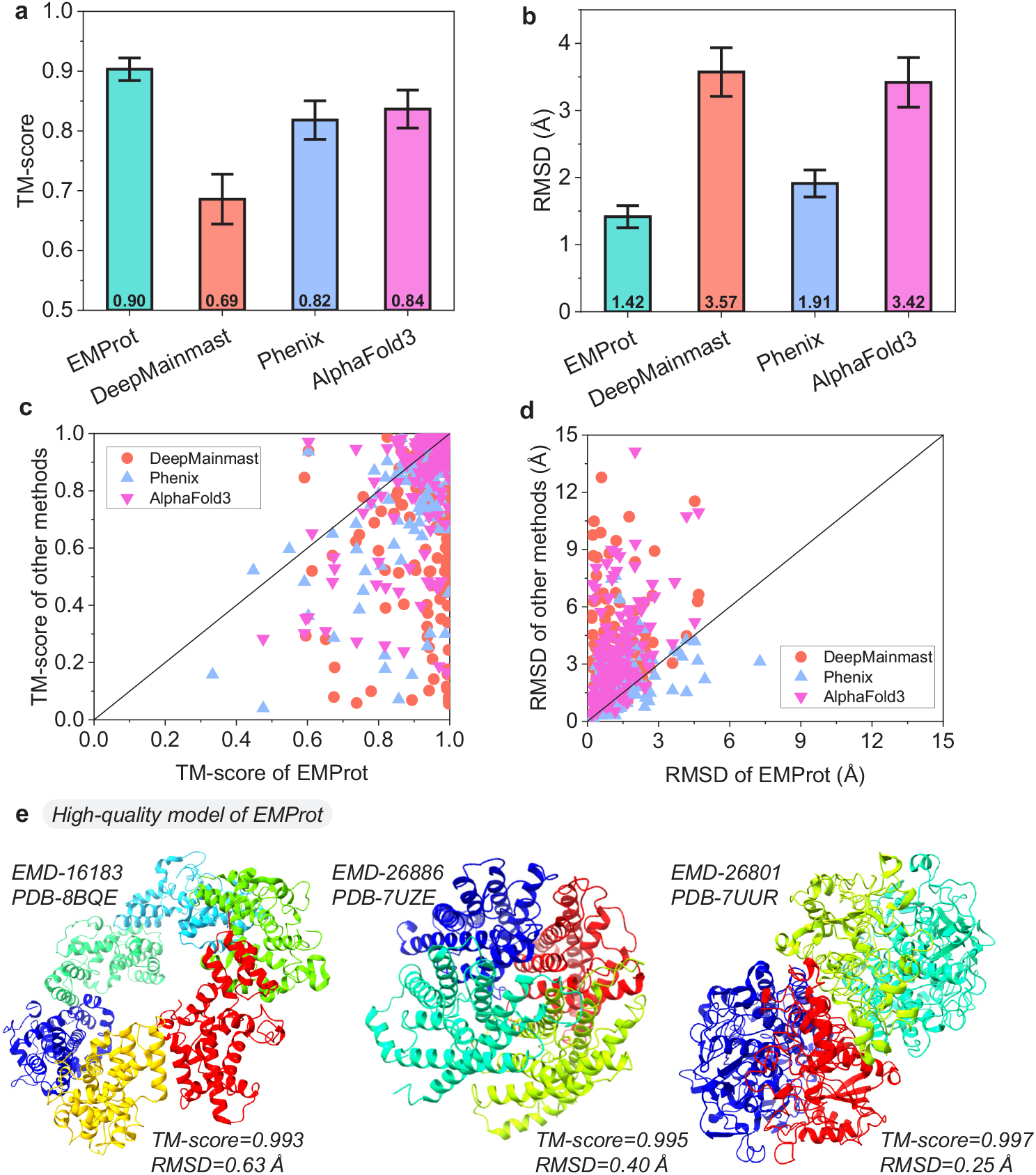
Comparison between EMProt, DeepMainmast, Phenix, and AF3 in building the complete structure on the test set of *n* = 177 maps. **a-b**, Bar plots of TM-score and RMSD. The bars indicate the means, error bars indicate 95% confidence interval. **c-d**, Head-to-head comparison of TM-score and RMSD. **e**, Three examples of high-quality models built by EMProt for three targets.

For RMSD (Fig. 4b, d), EMProt also shows an improvement over the other approaches. The average RMSD is 1.42 Å for EMProt, compared with 3.57 Å for DeepMainmast, 1.91 Å for Phenix, and 3.42 Å for AF3. Specifically, EMProt obtains a lower RMSD than DeepMainmast for 140 cases, Phenix for 101 cases, and AF3 for 142 cases. Due to lack of experimental map information, AF3 fails to build the models in atomic precision, and Phenix outperforms AF3 in RMSD conversely. The RMSD and TM-score calculation is based on matching query residues with target PDB residues globally, and only the matched residue contributes to the RMSD. EMProt achieves the highest TM-score while maintaining the lowest RMSD, suggesting that it can obtain the correct fold at atomic-accuracy levels.

Figure 4e show three EMProt high quality models for targets EMD-16183 (PDB: 8BQE), EMD-26885 (PDB: 7UZE), and EMD-26801 (PDB: 7UUR). All three models have a high TM-score of *>*0.99 and a low RMSD of *<*1.0 Å. Overall, there are 128 EMProt models that have a TM-score *>*0.90 and 37 cases that have a TM-score of *>*0.99 in the test set.

### 2.5 Examples in building the complete structure

Several examples are displayed in Fig. 5 to illustrate how the built models by EMProt, DeepMainmast, Phenix and AF3 is globally similar to the PDB structure. The built models are colored by chain names. Figure 5a shows the example for target EMD-15690 (PDB: 8AW3). For chain 2 colored in green, all map-to-model methods can build the structure correctly. The major challenge on this target is to build chain 3. In this regard, EMProt successfully build both chains and thus achieves a high TM-score of 0.888 for the built model, while DeepMainmast and Phenix fail to build the correct structure for chain 3 on this target, and AF3 cannot find the correct global fold, resulting the lower TM-score.

**Figure 5:**
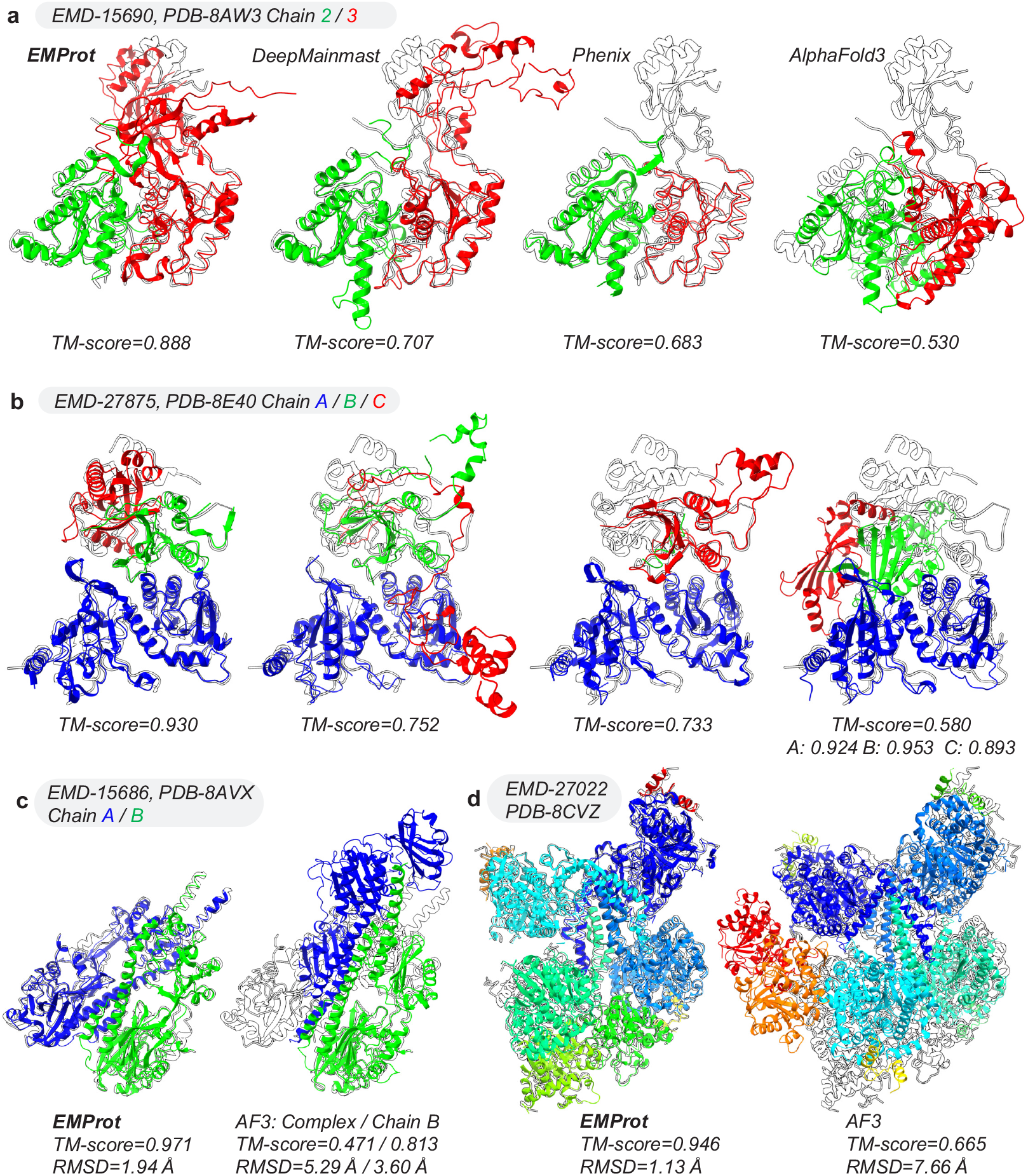
Comparison of the built models by EMProt, DeepMainmast, Phenix, and AF3 in terms of building the complete structures. **a**, Target EMD-15690 (PDB: 8AW3) at 3.6 Å resolution. **b**, Target EMD-27875 (PDB: 8E40) at 3.6 Å resolution. In **a, b**, nucleic-acids in the PDB model and the fragments built into the nucleic-acid regions are not shown. **c-d**, Comparison between EMProt and AF3 on targets EMD-15686 (PDB: 8AVX) at 3.5 Å resolution and EMD-27022 (PDB: 8CVZ) at 3.5 Å resolution.

Figure 5b show another examples of the built models by different methods for target EMD-27875 (PDB: 8E40). This is a small target with only three chains and a total of 636 residues. Although this target has near-atomic resolution of 3.57 Å, its density map has a severe heterogeneity, As such, ModelAngelo only yields a low sequence recall of 29.56% for this target. The major challenge for EMProt, DeepMainmast and Phenix on this target is to recover chain C in the model. Although EMProt achieves the best TM-score of 0.930, but one of the modeled chains (colored in red) still have a small deviation from the PDB model. DeepMainmast and Phenix both fails to recover a complete model and performs moderately on this target. Although AF3 predicts individual A/B/C chains well with high TM-scores of 0.924/0.953/0.893, the entire 3-chain complex is wrongly assembled with a low TM-score of 0.580, indicating the necessity of incorporating experimental map for accurate structure assembly.

We further compare EMProt and AF3 on two additional targets. Figure 5c shows the result of target EMD-15686 (PDB: 8AVX), a homologous dimer complex with a total of 961 residues. On this target, EMProt achieves a TM-score of 0.971 and an RMSD of 1.94 Å, which are much better than 0.471 and 5.29 Å for AF3. With the inclusion of experimental density map information, the complex is readily solved by EMProt with relatively high quality. However, the contacting interface is wrongly predicted by AF3, resulting a globally inverted conformation for chain A, though the monomer TM-score is still good (0.813). Figure 5d shows the example of another target, EMD-27022 (PDB: 8CVZ), including 10 chains and 3070 residues. The TM-score and RMSD are 0.946 and 1.13 Å for EMProt, which is much better than 0.665 and 7.66 Å for AF3. Further examination of the AF3 structure reveals that although the monomer prediction shows a high accuracy, there are still challenges in predicting protein multimer structures. These results suggest that without considering the experimental map information, methods like AF3 still face problems in modeling the correct structural assembly, especially for large complexes.

### 2.6 Model-to-map fit validation

Besides recovering the more structure from the map, another critical criterion for evaluating the accuracy of a built model is the consistency between the model and the map. The model-to-map fit assessment examines how well the atomic model explains the experimental data, and represents another golden-standard for structure modeling from cryo-EM maps. Here, four metrics, including CC mask, CC box, CC peaks and CC volume, which are calculated using the Phenix tool phenix.map model cc^35^, are used to evaluate the consistency of the built model with the map.

For a comprehensive comparison, we divided the methods into two groups of *de novo modeling* and *integrated modeling* during the evaluations. For *de novo* modeling, both the refined and unrefined models are evaluated. Overall, EMProt performs slightly better than ModelAngelo in *de novo* modeling (Table 1). Specifically, for the unrefined models, although EMProt yields slightly lower CC_mask and CC_volume values than ModelAngelo (0.700 and 0.704 versus 0.706 and 0.706), EMProt is better than ModelAngelo in terms of CC_box and CC_peaks (0.632 and 0.556 versus 0.627 and 0.550). After the model refinement, EMProt performs better than ModelAngelo in all four metrics and gives the CC_mask, CC_box, CC_peaks, and CC_volume values of 0.732, 0.656, 0.589, and 0.732, respectively, compared with 0.730, 0.647, 0.578, and 0.727 for ModelAngelo. Overall, the performance gaps between two methods for both model geometry and model-to-map fit are marginal, indicating the comparable performance of both methods.

**Table 1:**
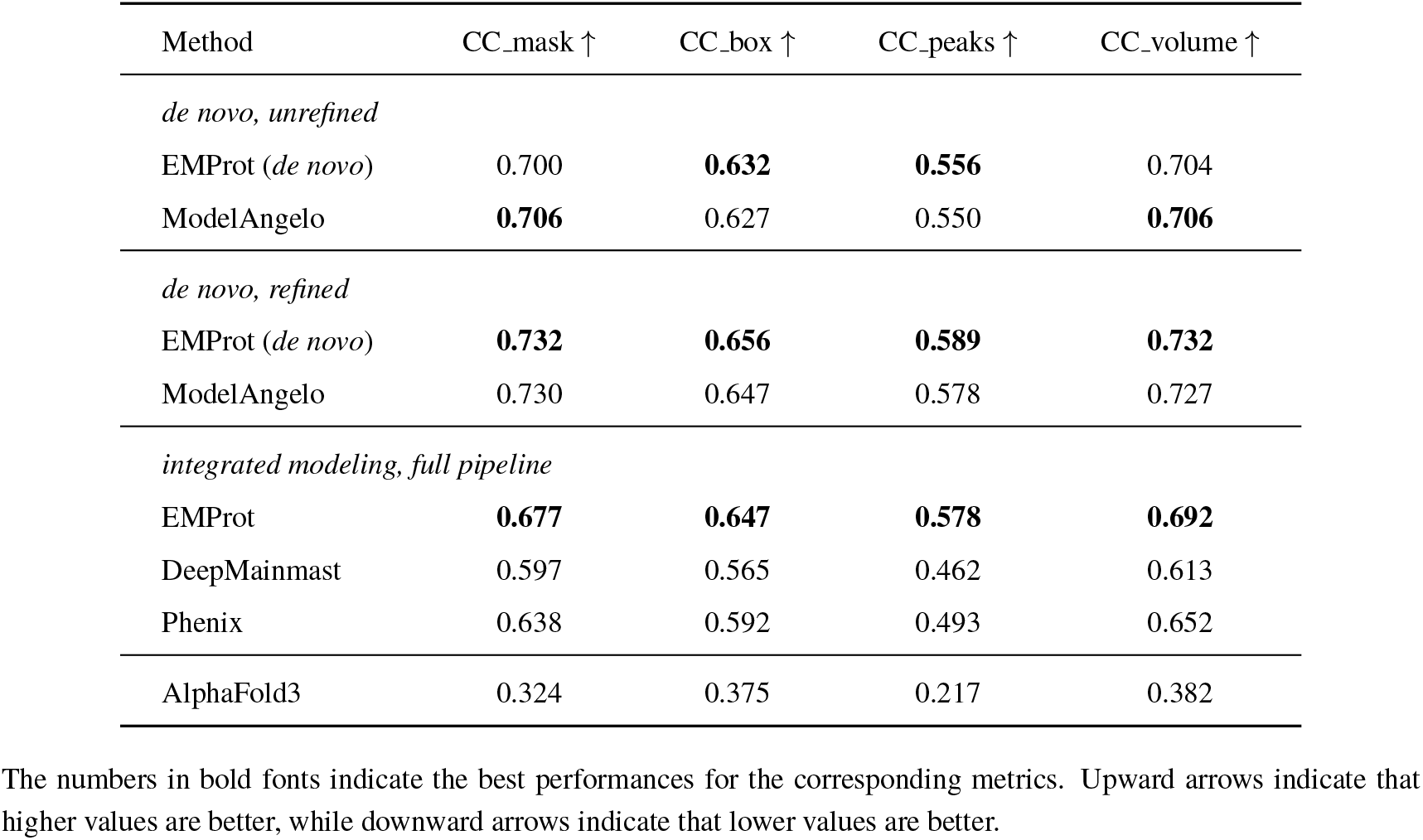
Model-to-map fit validation for the built models by EMProt, ModelAnglelo, DeepMainmast, and phenix.dock_and_rebuild (Phenix) on the test set of 177 cryo-EM maps. For reference, the results of the AF3 models are also listed.

| Method | CC_mask ↑ | CC_box ↑ | CC_peaks ↑ | CC_volume ↑ |
| --- | --- | --- | --- | --- |
| *de novo, unrefined* |  |  |  |  |
| EMProt (*de novo*) | 0.700 | **0.632** | **0.556** | 0.704 |
| ModelAngelo | **0.706** | 0.627 | 0.550 | **0.706** |
| *de novo, refined* |  |  |  |  |
| EMProt (*de novo*) | **0.732** | **0.656** | **0.589** | **0.732** |
| ModelAngelo | 0.730 | 0.647 | 0.578 | 0.727 |
| *integrated modeling, full pipeline* |  |  |  |  |
| EMProt | **0.677** | **0.647** | **0.578** | **0.692** |
| DeepMainmast | 0.597 | 0.565 | 0.462 | 0.613 |
| Phenix | 0.638 | 0.592 | 0.493 | 0.652 |
| AlphaFold3 | 0.324 | 0.375 | 0.217 | 0.382 |

The numbers in bold fonts indicate the best performances for the corresponding metrics. Upward arrows indicate that higher values are better, while downward arrows indicate that lower values are better.

When comparing the full-pipelines of EMProt and other methods, EMProt is consistently better than the other methods in all four metrics. Specifically, EMProt achieves the average CC_mask, CC_box, CC_peaks, and CC_volume values of 0.677, 0.647, 0.578, and 0.692, respectively, compared with 0.597, 0.565, 0.462, 0.613 for DeepMainmast, and 0.638, 0.592, 0.493, and 0.652 for Phenix (Table 1). It is noted that both EMProt and DeepMainmast adopts an integrated modeling strategy, but the model-to-map fit of EMProt is much better than DeepMainmast, indicating more effectively structure information integration of EMProt than DeepMainmast. In addition, the significantly better performance of EMProt than Phenix also indicates that fitting predicted models into cryo-EM maps is not trivial and simply docking and rebuilding predicted models into densities will not solve the modeling problem. As a reference, Table 1 also lists the results of original AF3-predicted models. The worst correlations between the AF3 models and the density maps again suggest the challenge and necessity of integrated model building for cryo-EM maps.

### 2.7 Structure validation of the build models

We next conduct all-atom structure vildation of the built models. Structure validation ensures that the built model has the reasonable geometric constraints, such as bond lengths, bond angles, torsion angles, and atom clash. Specifically, we use seven metrics to validate the build models, including Clashscore, MolProbity score, Ramachandran favored/outlier, Rotamer outlier, RMS (bonds) and RMS (angles), which are calculated by the Phenix tool phenix.molprobity^5, 36^. Along with the model-to-map fit metrics, these complementary evaluations provide a comprehensive measure of the model reliability and how close the models are ready to be deposited into the PDB.

Similarly, we first compare the performances between EMProt and ModelAngelo in *de novo* modeling. Overall, EMProt exhibits a comparable performance with ModelAngelo (Table 2). Specifically, for the unrefined models, although ModelAngelo is better than EMProt in terms of Clashscore, MolProbity score, and Rotamer outlier (66.51, 3.44, and 8.99% versus 69.36, 3.52, and 11.87%), EMProt is better than ModelAngelo in terms of Ramanchandran favored, Ramanchandran outlier, RMS (bonds), and RMS (angles) (92.90%, 2.12%, 0.0270, and 2.30 versus 91.80%, 2.78%, 0.0632, and 5.39). It is thus speculated that EMProt is better at modeling protein backbones, as measured by Ramanchandran facored/outlier, while ModelAngelo is better at modeling side-chains, as measured by Rotamer outlier. Moreover, EMProt is better at modeling standard amino-acid conformations, as measured by RMS (bonds) and RMS (angles), while ModelAngelo is better at modeling overall atom compatibility, as measured by Clashscore and MolProbity score. For the refined models, the performance differences between EMProt and ModelAngelo become smaller and more comparable (Table 2).

**Table 2:** Structure validation for the built models by EMProt, ModelAnglelo, DeepMainmast, and phenix.dock_and_rebuild (Phenix) on the test set of 177 cryo-EM maps. For reference, the results of the AF3 models are also listed.

| Method | Clash<br>score ↓ | MolProbity<br>score ↓ | Ramachandran<br>favored (%) ↑ | Ramachandran<br>outlier (%) ↓ | Rotamer<br>outlier (%) ↓ | RMS<br>(bonds) ↓ | RMS<br>(angles) ↓ |
| --- | --- | --- | --- | --- | --- | --- | --- |
| *de novo, unrefined* |  |  |  |  |  |  |  |
| EMProt (*de novo*) | 69.36 | 3.52 | **92.90** | **2.12** | 11.87 | **0.0270** | **2.30** |
| ModelAngelo | **66.51** | **3.44** | 91.80 | 2.78 | **8.99** | 0.0632 | 5.39 |
| *de novo, refined* |  |  |  |  |  |  |  |
| EMProt (*de novo*) | 10.41 | 1.99 | 95.59 | 0.29 | 2.54 | **0.0047** | **0.76** |
| ModelAngelo | **10.25** | **1.93** | **96.10** | **0.21** | **2.04** | 0.0098 | 0.87 |
| *integrated modeling, full pipeline* |  |  |  |  |  |  |  |
| EMProt | **13.44** | **2.01** | **95.92** | **0.26** | 1.97 | **0.0063** | **0.80** |
| DeepMainmast | 14.47 | 2.28 | 91.08 | 0.69 | **1.70** | 0.0080 | 0.96 |
| Phenix | 15.19 | 2.02 | 94.97 | 0.31 | 2.10 | 0.0068 | 0.83 |
| AlphaFold3 | 16.22 | 1.67 | 97.23 | 0.40 | 0.83 | 0.0112 | 1.49 |

The numbers in bold fonts indicate the best performances for the corresponding metrics. Upward arrows indicate that higher values are better, while downward arrows indicate that lower values are better.

We also evaluate the full-pipeline of EMProt, DeepMainmast and Phenix using the same metrics. As shown in Table 2, EMProt achieves an overall better performance than the other methods. Specifically, EMProt achieves a better performance than the other methods in six seven metrics, and yields the values of 13.44, 2.01, 95.92%, 0.26%, 0.0063, and 0.80 for Clashscore, MolProbity score, Ramachandran favored, Ramachandran outlier, RMS (bonds), and RMS (angles), respectively (Table 2). EMProt only performs worse than DeepMainmast in Rotamer outlier and still gives a good value of 1.97%, compared with 1.70% for DeepMainmast and 2.10% for Phenix. These results suggest the high accuracy of the built models by the full EMProt pipeline in terms of structure quality.

### 2.8 Quality assessment by local confidence scores

To assess the quality of the built model in a reference structure-free manner, we propose a local confidence score for each built residue. Specifically, we choose C*α* as the representative atom of a residue and define the local confidence score as the probability of the nearest voxel in the C*α* probability map. The local confidence score solely reflects the accuracy of C*α* atom placement, without considering the amino acid identity. We further define the displacement of a built residue as the distance between its C*α* atom and the nearest C*α* atom in the reference PDB structure. The confidence scores are grouped into bins of 0.05. As shown in Fig. 6a, the local confidence exhibits a good correlation with local displacement, suggesting that the local confidence score can effectively estimate the modeling accuracy of the built model at the residue level.

**Figure 6:**
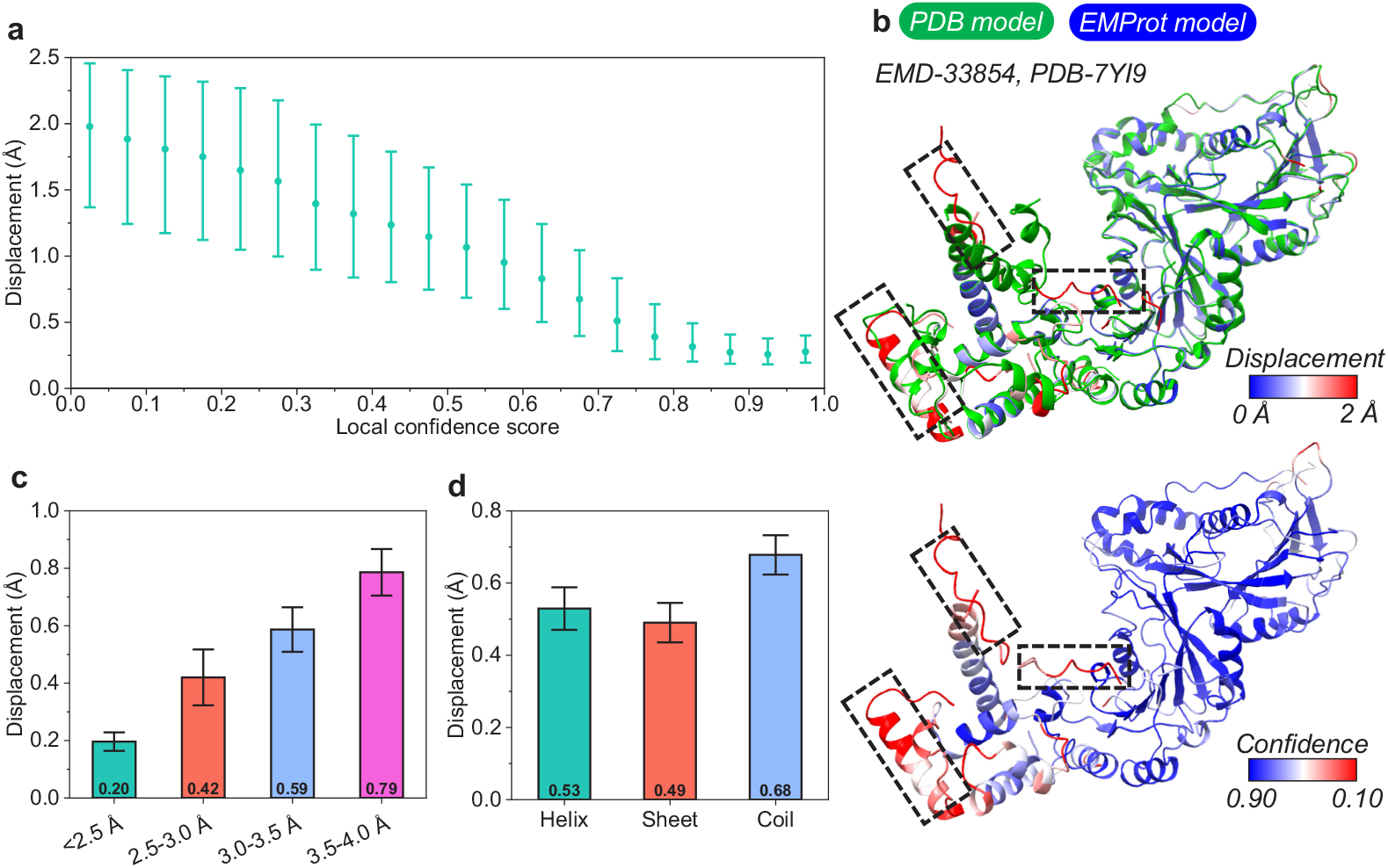
Quality assessment of the built models by local confidence scores. **a**, The residue displacements versus their local confidence scores for *n* = 371, 706 C*α* atoms that represent the corresponding residues on the test set of *n* = 177 maps. Confidences are merged into bins of 0.05. The dots indicate medians, and the error bars indicate first and third quartiles. **b**, An example of local confidence score as a displacement measurement, where the PDB model is colored in green and the EMProt model is in red-white-blue gradient. **c**, Bar plots of residue displacements for different global resolution bins. **d**, Bar plots of residue displacements for different secondary structure category. Error bars indicate 95% confidence interval.

One representative example, EMD-33807 (PDB: 7YG4), is shown in Fig. 6b. The PDB structure (colored in green) is superposed onto the EMProt-built model. In the upper panel, the built model is colored according to residue displacement, ranging from blue for 0 Å to red for 2 Å. The lower panel displays the same model, but the model is colored by local confidence score, from blue for 0.90 to red for 0.10. As shown in the figure, most residues are in blue in the built model, indicating relatively high modeling confidence and small displacement. However, some residues, particularly in coil/loop regions, exhibit larger deviations from the PDB structure, as indicated by dashed rectangles. Therefore, the proposed local confidence score is consistent with the residue displacement, and may be used to effectively estimate the modeling accuracy in a reference structure-free manner. However, a low confidence score does not necessarily indicate incorrect or highly deviated models, and may require a final round of expert inspection or manual pruning.

We further investigate the relationship between residue displacements and global resolutions in the EMProt-built models. Global resolutions are grouped into four bins: *<*2.5 Å, 2.5–3.0 Å, 3.0–3.5 Å, 3.5–4.0 Å. The bar-plots of the results are shown in Fig. 6c. It can be seen from the figure that EMProt achieves a good accuracy of *<*1.0 Å for all ranges of resolutions. Specifically, the average displacement for different resolution ranges are 0.20 Å, 0.42 Å, 0.59 Å, and 0.79 Å displacements increases as the resolution worsens, as expected. respectively. The

We also analyze the residue displacements in the built models for different secondary structure categories. The built residues are split into ‘Helix’, ‘Sheet’, and ‘Coil’ using secondary structure algorithm DSSP^37^. The bar-plots of the results are shown in Fig. 6d. Again, EMProt achieves a good accuracy of *<*1.0 Å for different structure categories. The average displacement for ‘Helix’ (0.53 Å) is worse than for ‘Sheet’ (0.49 Å), but both are better than for ‘Coil’ (0.68 Å). This phenomenon can be understood because helices and sheets exhibit more specific and stable spatial arrangements, while coils are more flexible. Additionally, coils are more prone to be erroneously modeled due to their lower resolutions in experimental maps compared to helices and sheets. Given the observation, it is recommended that users do manual inspection and modeling on coils more than helices and sheets.

## 3 Discussion

We have proposed an integrative modeling framework for accurate modeling of the complete protein structure from cryo-EM density maps, which referred to as EMProt. EMProt accurately builds the complete structure from cryo-EM maps through the integration of multiple well-designed submodules, including 3D recognition networks for main-chain atom prediction and residue type prediction, three-track-attention based structure refinement networks for all-atom construction, a fragment threading module for initial model building, and a fragment-fitting module for resolving missing residues that are hard to interpret from raw density maps.

EMProt is extensively evaluated on a benchmark set of 177 experimental cryo-EM maps and compared with state-of-the-art methods, including DeepMainmast, ModelAngelo, Phenix, and AlphaFold3. Overall, EMProt achieves the highest sequence recall in recovering the protein structure when compared to DeepMainmast, ModelAngelo, and Phenix, and yields the best TM-score and RMSD in building the complete structure when compared to DeepMainmast, Phenix and AF3. A key factor contributing to EMProt’s success lies in its effective integration of density-based *de novo* modeling with structure prediction. As the inclusion of predicted models in cryo-EM structure determination becomes increasingly mainstream, EMProt offers a practical and efficient solution for protein structure modeling from cryo-EM maps.

Through the comprehensive evaluation and comparison with related methods, EMProt exhibits improved performances over state-of-the-art methods. On one hand, EMProt can *de novo* model protein structures from cryo-EM density maps, achieving a performance comparable to the leading *de novo* method, ModelAngelo. On the other hand, by incorporating AF2 structures, EMProt significantly improves the completeness of the built models while maintaining overall good model geometry and model-map agreement. Nevertheless, the resulting models are not sufficiently perfect to omit subsequent manual processing steps. Therefore, it is recommended that users refine the built models with a third-party program like phenix.real space refine before further analysis. In summary, EMProt marks a major step forward in cryo-EM structure modeling. Future improvements could further enhance model quality, particularly for difficult-to-resolve regions, paving the way for even more reliable and high-throughput protein structure determination.

## 4 Methods

### 4.1 Data collection

The test set used in this study consists of 177 non-redundant cryo-EM maps at *<* 4 Å resolutions sourced from the ModelAngelo paper^17^. The test set is constructed by excluding the targets that (i) have *>*30,000 residues, (ii) are viruses with I-group symmetry, and (iii) contain insertion codes or other irregularities.

To construct a training set for the neural networks, we first queried the EMDB for a complete list of maps with resolutions *<* 4 Å and their corresponding PDB structures ^9, 38^. To remove redundancy between the test and training sets, we adopted MMseqs2^39^ to search for redundant chains in the PDB structures, using a sequence identity cutoff of 30%. If any chain in a candidate PDB structure is similar to a chain in the test set, the associated map and PDB structure are removed. The remaining maps and PDB structures are further filtered based on the following criteria: (i) The maps with non-orthogonal axes are excluded; (ii) The maps reconstructed by methods other than single-particle analysis are discarded; (iii) The PDB structures with unassigned amino acid types or containing only backbone atoms are removed. Next, we calculate the cross-correlations between the experimental maps and the structure-simulated maps using Chimera^40^. The map-structure pairs with cross-correlations below a threshold of 0.60, indicating severe misfits, are filtered out. The remaining maps are then manually inspected for quality control. Subsequently, we apply MMseqs2 to cluster the chains with a sequence identity cutoff of 30%. To further reduce the redundancy, we develop an in-house greedy algorithm to cluster the corresponding PDB items based on the MMseqs2 clustering results. Specifically, if two chains from different PDB items are identified as similar, only one of the PDB items is retained. Finally, a total of 802 maps and their corresponding PDB structures are retained as the training set.

### 4.2 Main-chain atom and residue type prediction from density map

#### 4.2.1 The Swin-Conv-UNet network architecture

We employ a Swin-Conv-UNet (SCUNet)^41^ architecture to extract atom probabilities and amino acid identities from density maps. The network consists of three encoder blocks, one transition block, and three decoder blocks, which are based on Swin-Conv (SC)^41, 42^ modules, with skip connections linking the encoders and decoders. Each Swin-Conv block contains a Swin transformer (SwinT) block parallelized with a residual convolutional (RConv) block, which is sandwiched between two 1*×*1 convolutions. The Swin transformer uses a window size of 3. For down-sampling, the network applies a 3D convolution layer with a kernel size and stride of 2. For up-sampling, it uses a 3D transposed convolution layer with the same kernel size and stride. The input to the network consists of density chunks of size 48 *×* 48 *×* 48 with a grid interval of 1.0 Å. The output dimensions are the same as the input. Both atom probability and amino acid identity predictions utilize the same network architecture, differing only in the number of output channels in the final layer.

#### 4.2.2 Network training

To train the main-chain atom probability predictor, the N, C*α*, and C atoms are extracted from the structure, and the training labels are generated based on the atom positions and map resolution. For each C*α* atom, the density value at grid point **x** is calculated as follows^43, 44^:

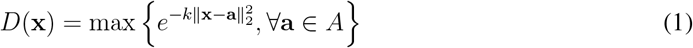

where **a** denotes the coordinate of C*α* atom in structure *A*, and the kernel *k* is defined according to the map resolution *R* as ^45^

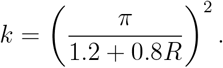

It should be noted that here a local resolution might be used to get a more precise representation of local density quality. However, accurate local resolution requires the calculation between two half maps, which may not be achievable because the authors do not always deposit the half maps into the EMDB. Thus, we used a global resolution in this study for simplicity.

The N and C atom probability maps are generated in the same manner and concatenated with the C*α* probability maps. The loss for atom probability prediction is a combination of Smooth L1 loss and structural similarity index measure (SSIM)^46, 47^ loss, as follows^21^:

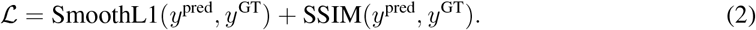

To train the amino acid type segmentation network, each voxel is labeled with the nearest amino acid type [0, 20) within 3 Å. Voxels that do not correspond to any amino acid within this distance are labeled as background. The loss function is a 20-class cross-entropy loss, with label-20 predictions ignored, as only predictions near the backbone are of interest:

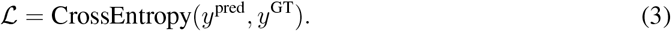

To process the training data, the grid size of the map is standardized to 1.0 Å using trilinear interpolation. The density values of each experimental map are clipped to the range between ≥ 0.0 and the ≤ 99.999% percentile of the density values, and then normalized to [0, 1]. The experimental maps and labeled maps are partitioned into overlapping chunks of size 60 *×* 60 *×* 60 with a stride of 30. For computational efficiency, those experimental chunks with a maximum density value less than 0.0 are excluded from training.

During training, data augmentation is applied to prevent overfitting. The input box is randomly cropped to 48 *×* 48 *×* 48 from the 60 *×* 60 *×* 60 chunk, and a random 90° rotation is applied. The network is implemented using PyTorch 2.1. The Adam optimizer is used to update the network weights. We train the atom probability predictor and the amino acid type predictor separately using the same hyperparameters. The initial learning rate is set to 5 *×* 10^*−*4^, with an L2 regularization coefficient (weight decay) of 1 *×* 10^*−*4^. During training, the learning rate is halved if the average training loss does not decrease for 4 consecutive epochs, with a minimum learning rate capped at 5 *×* 10^*−*6^. Each network is trained with a batch size of 96 on two Nvidia A100 GPUs (40 GB). The models with the lowest validation loss are selected as the final models.

### 4.3 All-atom construction in density map

#### 4.3.1 Three-track attention network architecture

After getting the C*α* atom probability, the C*α* atoms are derived from the probability map by detecting local maxima using a mean-shift algorithm. We then use a three-track-attention network to predict backbone frames (*R*, **t**) and torsion angles *τ* for each C*α* atom. It consists of 16 main blocks, iteratively processing 1D, 2D, and 3D information.

The densities around each C*α* (a cubic with shape of 23^3^ Å ^3^) are extracted as initial 1D features. The densities between pairwise C*α* atoms (a cuboid with shape of 17*×*3*×*3 Å ^3^) are extracted as initial 2D features. For information flowing from 1D, 2D, and 3D to 1D, we used self-attention and invariant point attention module to update 1D features, with attention biases from 2D features and 3D frames. For information flowing from 1D to 2D, we used outer product to update the 2D features. For information flowing from 2D and 3D to 2D, we used axial-attention to update the 2D features, with attention biases from 3D frames. The backbone update module predicted the new backbone frames for each C*α* atom, giving the updated 1D features. After going through 16 blocks, a final layer is used to predict torsion angles. Having the backbone frames and torsion angles, all-atom positions can be determined through a three-track attention network.

#### 4.3.2 Network training

The neural network is trained using a customized noising/denoising schedule. For each residue, its C*α* position **t**^*r*^ is randomly noised according to the following formula:

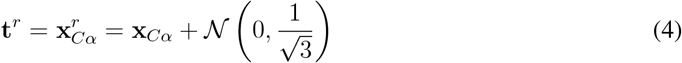

where 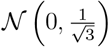 is a random gaussian noise applied to the C*α* position. The rotation matrix *R*^*r*^ of the backbone frame is noised using the following procedure:

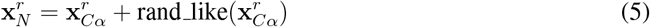

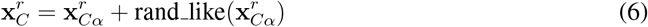

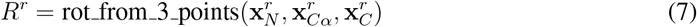

The training procedure consists of the following steps: (1) selecting a PDB structure, (2) randomly choosing a residue, (3) extracting its *k* neighboring residues (including itself), (4) applying noise to each residue, and (5) using the neural network to predict the true positions. For training, we use an effective batch size of 8 and set the number of neighbors to *k* = 128. The neural network parameters are updated using AdamW optimizer. The initial learning rate is set to 1 *×* 10^*−*4^. We stop the gradient of rotation between consecutive blocks to stabilize the training.

The loss function consists of three learning objectives. The first loss is the RMSD loss, which is defined on the C*α* atoms, backbone N, C*α*, and C atoms, and all atoms. C*α* and backbone RMSD losses are calculated in every intermediate layer of the network, with the *l*-th layer having a weight of *γ*^16*−l*^ (*γ* is set to 0.80 in this study). The C*α* RMSD loss is defined as

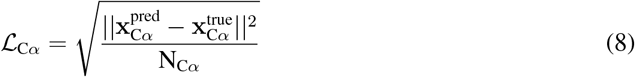

and the backbone/all-atom loss is defined for backbones and all-atoms similarly. The total RMSD loss is defined as

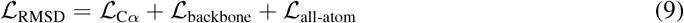

The second loss is the torsion angle loss, which is defined as

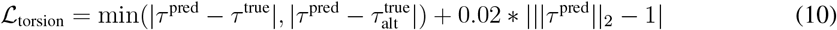

The third loss is the structure violation loss *ℒ*_violation_. The structure violation loss penalize the structure irregularity, e.g. atom clash, irregular bond-lengths and bond-angles. The total loss for network training is the sum of all three losses as follows.

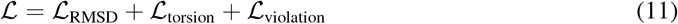

### 4.4 Backbone tracing

Since the predicted all-atom residues are unordered, it is needed to connect residues to chains. A peptide-bond 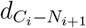 is defined as the distance between C atom in the last residue and N atom in the next residue. Given the fact that 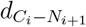 only varies in small ranges, if we start from one residue, we can iteratively find its nearest next residue under the *C − N* distance restraint (*d <* 2.1) and the residues are threaded to fragments.

### 4.5 Sequence assignment

For each residue in a fragment, we locate the nearest voxel to its C*α* atom in the predicted amino-acid type map, and assigns the predicted logits to the residue. For each fragment, the amino-acid logits are used to construct an HMM profile. A sequence search is then performed against the given sequences using HMMER. After the HMM search, residues are assigned with the corresponding amino acid types according to the alignment results.

### 4.6 Integration of predicted structures

#### 4.6.1 Structure prediction

For each chain, the full-length sequence in the PDB SEQRES record is used as the input to AF2 (v2.3.0). The max-template-date is set to 2022-05-01, a time point before all the test models are released. We truncate the predicted model to have the same start and end residue using a simple sequence alignment. It is noted that 22 chains in eight PDB structures failed in the AF2 prediction because their too long sequences, for which we used AF3^27^ to predict their structures instead.

#### 4.6.2 Chain partition

The predicted protein models, although of high accuracy, may have large deviation in inter-fragment level. To find the best match with the map, split of chains into sub-components is necessary. Following UniDoc^48^, chains are split to domain level and fragment level based on structure contacts. The C*α* distances are converted to contact map *p*_*m,n*_, which is then used to calculate the inter-domain interactions score S_inter_(*D*_*i*_, *D*_*k*_) and intra-domain interaction score S_intra_(*D*_*i*_). The definition of these scores can be found in the UniDoc study. In a top-down splitting stage, the structure is first cut into fragments to minimize S_inter_ iteratively. In a bottom-up merging stage, the fragments are merged to maximize the score S(*i, k*) iteratively

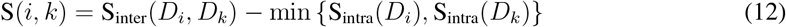

With such splitting-merging strategy, a chain can be split to fragments (before merging) and domains (after merging). A chain contains *>*= 1 domain(s), and a domain contains *>*= 1 fragment(s).

#### 4.6.3 Matching template by structure alignment

To combine the information from the template structure (AF2 models in this study) and the density map, one way is to align the template to the *de novo* chains directly built from the map. The chains built from the map provides a highly accurate backbone information but the predicted AlphaFold models may have large movement between domains^49^. Therefore, the first step is to split each template structure into sub-structure and perform the matching at sub-structure level. We split the template into fragments, and the matching is performed on fragments.

Each sub-structure is aligned to the built chains using TMalign^32^. If the RMSD between the aligned residues is *<* 1.5 Å, the sub-structure is considered as being matched. The coordinates of the matched sub-structure are then transformed using the output transformation matrix from TMalign. The matched sub-structures also helps to correct the wrong sequence alignment and fix missing residues on the built chains. If the aligned residues between the sub-structure and the chain has a sequence identity of *>* 80%, which means they are highly matched in sequence, then the sequence of the matched residues is updated to the corresponding sequence in the sub-structure and the missing residues are fixed using AF2 residues.

#### 4.6.4 Matching template by structure fitting

Besides aligning the template structure to the built chains, EMProt also directly fit the template structure to the density map. Especially when local resolution are poorer, *de novo* modeling often fails to give a chain model but template fitting fills the coverage gap. Rather than fitting to the raw map, we fit the template to the predicted main-chain map since it is less noised.

EMProt uses an exhaustive search in rotational and translation space to maximize the matching score between structure and main-chain map:

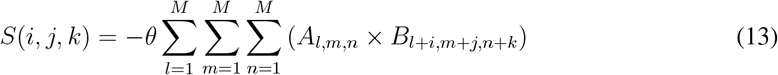

where grid A is generated by mapping the N, C*α*, and C atoms of the structure into a 3D grid, and grid B denotes the main-chain map. Grid A is sampled with a grid interval of 3.0 Å and grid B is down-sampled from the main-chain map to have the same grid interval. An angle interval of 18° is used to evenly discretize the Euler space, resulting in a total of 2540 evenly distributed euler angles. The searching process is accelerated by fast-Fourier transform (FFT) and multi-thread programming. After clustering the sampled poses with an RMSD cutoff of 5 Å, each structure is placed at most *M* different positions.

When fitting structure to the map, a semi-flexible refinement is performed to improve local sub-structure conformation. We employ a local SIMPLEX optimization algorithm to find the best match of the sub-structures. For a template structure, starting from a selected sub-structure, the positions of all sub-structures are optimized one after another though a breadth-first search (BFS) on the graph defined by sub-structure adjacency. Taking each sub-structure as a seed, the refinement will output *n* different structure models. The results of sub-structure refinement for one structure are *M × n* refined models plus *M* rigidly fitted models.

We adopted two strategy for fitting structures to the map. First, we fit the structure of the chains, where the sub-structure are the domains. Second, we fit the structure of the domain, where the sub-structure are the fragments. For both strategy, all chains/domains are split into fragments for finial assembling.

#### 4.6.5 Structure assembling from fragments

After the matching step by structure alignment and fitting, the fitted AF2 fragments and the fixed built chains forms a structure library. These fragments are gathered for a final stage of assembling. For each fragment, a match score is defined as

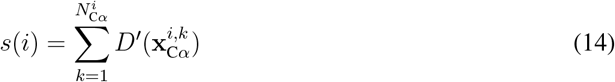

where *D*^*′*^(**x**) is the predicted atom probability map (the density values are normalized between 0.0 and 99.999 percentile value). For fragments *i* and *k*, if the distance between a C*α* atom in fragment *i* and another C*α* atom in fragment *k* is *<* 2.0 Å, the two atoms are clashed and the clash percentage of fragment *i* is defined as

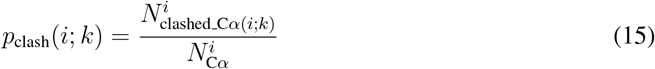

where 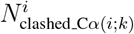is the number of clashed C*α* atoms and 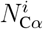 is the total C*α* atoms in fragment *I* and *p*_clash_(*k*; *i*) is defined in the same way. Two fragments *i* and *k* are clashed if *p*_clash_(*i*; *k*) *> p*_cutoff_ or *p*_clash_(*k*; *i*) *> p*_cutoff_. The problem of assembling the fragments is then converted to a constrained optimization problem, formulated as maximizing

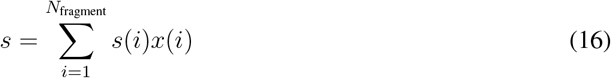

subject to

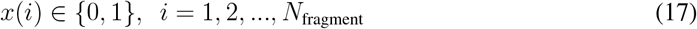

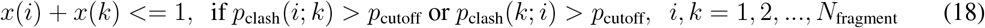

We set *p*_cutoff_ to 0.10 in this study, allowing a mild clash between fragments. Generally, if two fragments are clashed, we expect to keep the larger one. The assembled fragments are input to the all-atom refinement network again for a final round of all-atom refinement.

### 4.7 Post refinement

After the structure is assembled, users can refine the built model to improve local structure and model-to-map fit through a third-party program, which is highly recommended in real applications. For a fair evaluation and comparison, in this study, the built models by all methods, including EMProt, ModelAngelo, DeepMainmast, and Phenix, are refined using phenix.real space refine^33^ with the default parameters, unless otherwise specified.

### 4.8 Evaluation of the built models

Two types of evaluations are used to assess the accuracy of a built model using the PDB structure as the reference. One is recovering the protein structure, that is to measure how much the PDB structure is recovered by the built model regardless of the order and connectivity of built fragments. The other is building the complete structure, that is to measure how much the built model is similar to the PDB structure in a global way.

For recovering the protein structure, three metrics, including residue coverage, sequence match, and sequence recall, are used to measure the accuracy of a built protein model. The metrics are calculated using a distance cutoff of 3 Å and C*α* as the representative atom for each residue using phenix.chain comparion^10^. Here, the residue coverage measures the fraction of residues in the built model that lie within a specified distance cutoff from the map, irrespective of residue type, relative to the PDB structure. Sequence match indicates the fraction of the residues in the built model that have the same residue type as those in the PDB structure. The sequence recall is defined as the product of residue coverage and sequence match. It reflects the proportion of residues in the built model that are correctly placed in terms of both backbone position and residue type when compared to the PDB structure.

For building the complete structure, two metrics, TM-score and RMSD, are used to measure the global similarity between the built model and the PDB structure. The two metrics are calculated using the USalign program^34^.

In addition, we also use four other metrics, including CC_mask, CC_box, CC_peaks, and CC_volume calculated by phenix.map_model_cc^35^, to evaluate model-map consistency. It is noted that the CC values vary with model’s B-factors. Since the B-factors presented by different methods are in different numerical ranges, we reset all model’s B-factors to 0.0 for simplicity. For structure validation of the build models, we use seven metrics, including Clashscore, MolProbity score, Ramachandran favored/outlier, Rotamer outlier, RMS(bonds) and RMS(angles) calculated by phenix.molprobity^5, 36^.

## Data availability

The raw data of the evaluation results are provided in the Article. All published data sets used in this paper were taken from the EMDB and PDB.

## Code availability

The EMProt package is freely available for academic or non-commercial users at http://huanglab.phys.hust.edu.cn/EMProt.

## Acknowledgements

This work was supported by the National Natural Science Foundation of China (grant Nos. 32161133002, 32430020, and 62072199), the startup grant of Huazhong University of Science and Technology and the Postdoctoral Fellowship Program of CPSF (grant No. GZB20250617).

## Author contributions

S.H. conceived and supervised the project. T.L., J.C, H.L., and H.C. designed and performed the experiments. S.H. and T.L analyzed the data. T.L. and S.H. wrote the manuscript. All authors reviewed and approved the final version of the manuscript.

